# Sox2 and canonical Wnt signaling interact to activate a developmental checkpoint coordinating morphogenesis with mesodermal fate acquisition

**DOI:** 10.1101/2020.01.29.924050

**Authors:** Brian A. Kinney, Richard H. Row, Yu-Jung Tseng, Maxwell D. Weidmann, Holger Knaut, Benjamin L. Martin

## Abstract

Animal embryogenesis requires a precise coordination between morphogenesis and cell fate specification. It is unclear if there are mechanisms that prevent uncoupling of these processes to ensure robust development. During mesoderm induction, mesodermal fate acquisition is tightly coordinated with the morphogenetic process of epithelial to mesenchymal transition (EMT). In zebrafish, cells exist transiently in a partial EMT state during mesoderm induction. Here we show that cells expressing the neural inducing transcription factor Sox2 are held in the partial EMT state, stopping them from completing the EMT and joining the mesodermal territory. This is critical for preventing ectopic neural tissue from forming. The mechanism involves specific interactions between Sox2 and the mesoderm inducing canonical Wnt signaling pathway. When Wnt signaling is inhibited in Sox2 expressing cells trapped in the partial EMT, cells are now able to exit into the mesodermal territory, but form an ectopic spinal cord instead of mesoderm. Our work identifies a critical developmental checkpoint that ensures that morphogenetic movements establishing the mesodermal germ layer are accompanied by robust mesodermal cell fate acquisition.

## Introduction

Epithelial to mesenchymal transition (EMT) is the process in which epithelial cells lose their adhesion to neighboring cells and adopt a mesenchymal migratory phenotype. This process was first described by observing chick mesoderm formation (Hay 1995), and was later found to occur in many other normal processes, as well as disease states such as cancer metastasis (Nakaya and Sheng 2013; Nieto 2013). More recently, metastable partial (also referred to as intermediate) EMT states have been observed, where cells maintain a transitional state that shares characteristics of both epithelial and mesenchymal cells (Ye and Weinberg 2015; Li and Kang 2016; Nieto et al. 2016). Partial EMT states are thought to be particularly important in the process of solid tumor metastasis, where metastable partial EMT states exhibit increased migratory and invasive capacity, as well as more stem-cell like characters (Campbell 2018; Aiello and Kang 2019). Despite this, it is unclear what purpose, if any, metastable partial EMT states play during normal development.

Vertebrate embryos contain neuromesodermal progenitors (NMPs) (Kimelman 2016; Martin 2016), which make a binary decision to become spinal cord cells, or mesoderm that will primarily form the somites (Tzouanacou et al. 2009; Martin and Kimelman 2012). During mesoderm induction, NMPs undergo an EMT, which is tightly associated with the acquisition of mesodermal fate (Goto et al. 2017). This occurs in a two-step process, where Wnt signaling initiates the EMT, and FGF signaling promotes EMT completion by activating the expression of the transcription factors *tbx16* and *msgn1* (Goto et al. 2017). Zebrafish embryos deficient in the t-box transcription factor *tbx16* (originally called *spadetail*) have a large accumulation of cells at the posterior-most structure of the embryo called the tailbud (Kimmel et al. 1989; Griffin et al. 1998), a phenotype caused by the inability of the NMPs to complete their EMT and join the developing paraxial mesoderm (Row et al. 2011; Manning and Kimelman 2015). This phenotype is very similar to mouse embryos lacking function of the related t-box transcription factor *Tbx6*, which also have an enlarged tailbud and deficit in paraxial mesoderm (Chapman and Papaioannou 1998). Cells in the partial EMT state exhibit increased adhesiveness compared to the fully mesenchymal state, and cells lacking *tbx16* maintain a metastable partial EMT state until *tbx16* is activated, after which they complete the EMT (Row et al. 2011).

NMPs are characterized by co-expression of the two transcription factors, *sox2*, which promotes spinal cord fate, and *brachyury* (*tbxta* and *tbxtb* in zebrafish), which specifies mesoderm in part through activation of canonical Wnt signaling (Martin and Kimelman 2008; Martin and Kimelman 2010; Takemoto et al. 2011; Martin and Kimelman 2012; Bouldin et al. 2015). Here we show that the critical role of Tbx16 is to repress *sox2* transcription in the partial EMT state as cells become mesoderm. Sox2 activation alone is sufficient to recapitulate the Tbx16 loss of function phenotype, where cells are prevented from exiting the tailbud and remain trapped in an undifferentiated partial EMT state. This acts as a developmental checkpoint, since cells with *sox2* expression in mesodermal territories outside of the tailbud will become neurons. Thus, the checkpoint ensures coordination of morphogenesis with proper cell fate acquisition to prevent ectopic neural formation. Our work for the first time demonstrates an essential normal function of partial EMT states during development, and provides insight into how the partial EMT state in cancer can be targeted by inhibiting developmental checkpoints.

## Results

### Sox2 activation is sufficient to induce neural differentiation in a context dependent manner

NMPs express the neural inducing transcription factor *sox2*, which is down-regulated as NMPs become mesoderm (Delfino-Machin et al. 2005; Takemoto et al. 2011; Martin and Kimelman 2012; Bouldin et al. 2015). To determine the role that Sox2 plays in NMPs, we used a heat-shock inducible *sox2* transgenic line (*HS:sox2*) (Row et al. 2016). When *sox2* was activated throughout the embryo at the end of gastrulation (bud stage) and analyzed at 24 hours post fertilization (hpf), ectopic expression of the neural marker *neurog1* was observed in mesodermal territories (Fig. 1A, B), and there was a corresponding decrease in the skeletal muscle marker *myod* (Fig. 1C, D). Activation of *sox2* at bud stage in the background of reporter transgenes for skeletal muscle (*actc1b:gfp*) (Higashijima et al. 1997) and neurons (*neurog1:mkate2*) resulted in a loss of differentiated muscle and the presence of ectopic neurons in mesodermal territories (Fig. 1E, F). To test whether the effect of Sox2 is cell autonomous, we transplanted cells transgenic for *HS:sox2* and *neurog1:mkate2* (Fig. 1G-H’) or *HS:sox2* and *actc1b:gfp* (Fig. 1I-J’) into the ventral margin of shield stage wild-type embryos. The activation of *sox2* by heat shock in transplanted cells caused a significant cell-autonomous induction of more neural cells, including in ectopic locations, at the expense of skeletal muscle cells (Fig. 1G-L, for *neurog1:mkate2* quantification 1,177 wild-type donor cells were counted in 8 host embryos, and 2,051 *HS:sox2* donor cells were counted from 10 host embryos, statistics were performed using an unpaired t test, P=*0.0105, for *actc1b:gfp* quantification 1,307 wild-type donor cells were counted in 8 host embryos, and 971 *HS:sox2* donor cells were counted from 5 host embryos, statistics were performed using an unpaired t test ***P=0.0003). Intriguingly though, the induction of ectopic neural fate by *sox2* in both the whole embryo and transplant conditions was localized to more anterior regions of the embryo. Additionally, while 72.54% of control transplanted cells differentiated into either muscle or neurons, only 42% of *sox2*-expressing cells differentiated into these cell types (Fig. 1K, L). Upon further examination, a large proportion of *sox2*-expressing cells gave rise to fin mesenchyme, a phenotype that is also observed in transplanted *tbx16* mutant cells, which are trapped in the partial EMT state (Fig. 1M-P, 2,129 wild-type donor cells were counted in 6 host embryos, 2,714 tbx6 -/- donor cells were counted in 6 host embryos (***P=0.0006), and 2,347 *HS:sox2* donor cells were counted from 9 host embryos (*P=0.0322)) (Ho and Kane 1990; Row et al. 2011).

**Figure 1.**
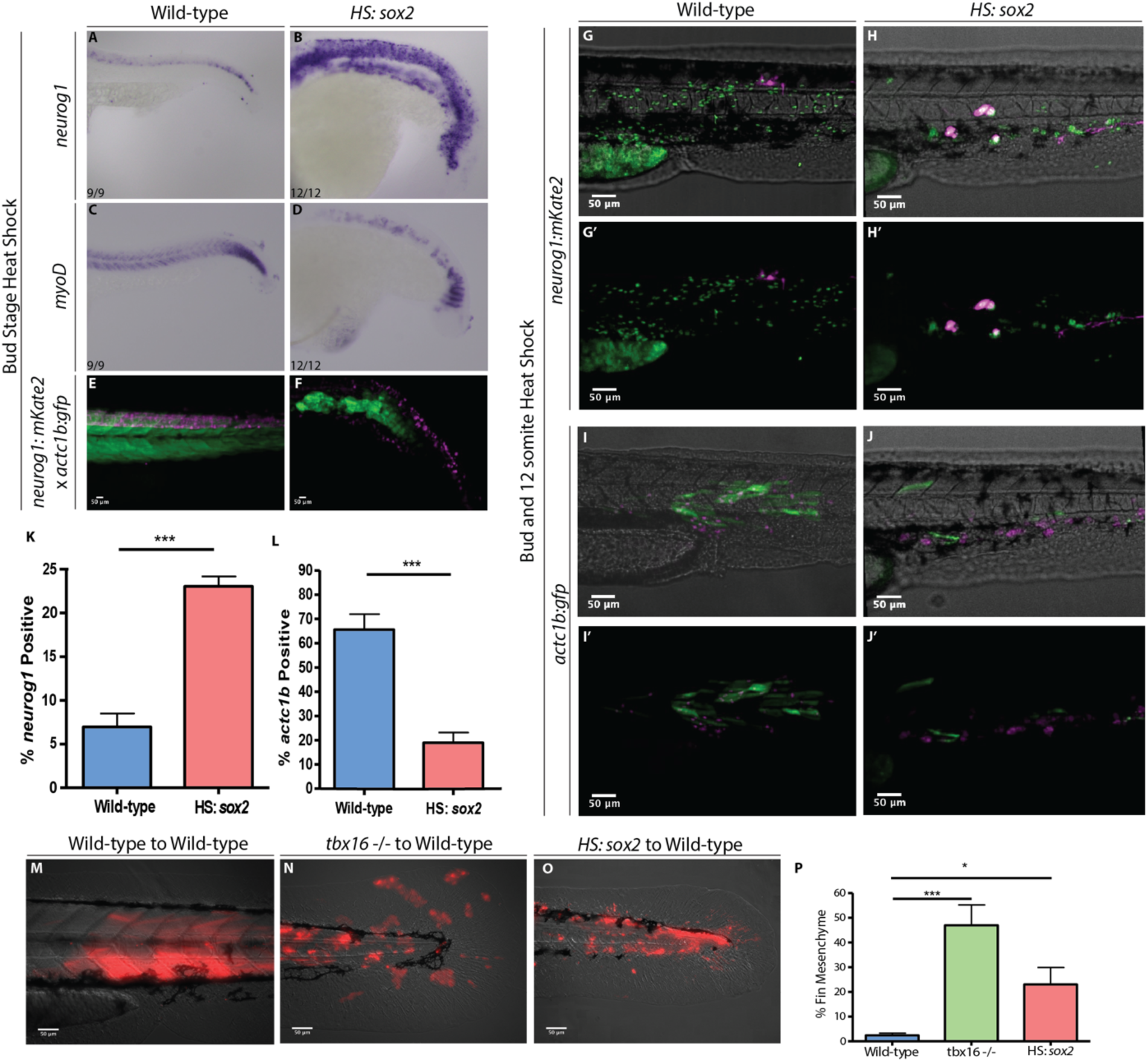
*sox2* activation causes an increase of neural progenitors and a decrease in presomitic mesoderm. Whole-mount in situ hybridization visualizing *neurog1* (neural) (A, B) or *myod* (skeletal muscle) (C, D) in wild-type (A, C) and *HS:sox2* embryos (B, D). All embryos for *in situ* hybridization were heat shocked at bud stage at 40°C for 30 minutes and fixed at 24 hpf. Transgenic embryos with the *ngn:mKate* and *actc1b:gfp* reporters show a similar neural expansion and muscle loss in *HS:sox2* (F) embryos compared to wild-type (E). Live-imaged transgenic embryos were heat-shocked at bud stage at 40°C for 30 minutes and imaged at 36 hpf. Embryos with the *neurog1:mkate* (G-H’) or the *actc1b:gfp* (I-J’) reporter were injected with *NLS-KikGR* mRNA and transplanted into the ventral margin of wild-type host embryos. Donor cells with the *HS:sox2* transgene exhibited an increase in the percentage *neurog1:mkate* positive cells (H, H’ compared to G, G’ and quantified in K, 1,177 wild-type donor cells were counted in 8 host embryos, and 2,051 *HS:sox2* donor cells were counted from 10 host embryos, statistics were performed using an unpaired t test, P=*0.0105) and a decrease in the percentage of *actc1b:gfp* positive cells (J, J’ compared to I, I’ and quantified in L, 1,307 wild-type donor cells were counted in 8 host embryos, and 971 *HS:sox2* donor cells were counted from 5 host embryos, statistics were performed using an unpaired t test ***P=0.0003). The NLS-KikGR protein was photoconverted to red fluorescence in I-J’. Wild-type, *tbx16* mutant, and *HS:sox2* embryos were injected with rhodamine dextran and transplanted into the ventral margin of shield stage wild-type host embryos (M-O). The percent of transplanted cell contribution to fin mesenchyme is quantified in panel P (2,129 wild-type donor cells were counted in 6 host embryos, 2,714 tbx6 -/- donor cells were counted in 6 host embryos (***P=0.0006), and 2,347 *HS:sox2* donor cells were counted from 9 host embryos (*P=0.0322)). All transplants were heat shocked at bud stage and 12-somites at 39°C for 30 minutes.

### Sustained *sox2* expression in mesoderm fated NMPs traps them in a partial EMT state

Tbx16 is necessary and sufficient for *sox2* repression (Bouldin et al. 2015), and *tbx16* loss-of-function and *sox2* gain-of-function both bias transplanted cells located in the tailbud of host embryos towards a fin mesenchyme fate (Fig. 1M-P). We hypothesize based on these observations that maintenance of *sox2* expression in *tbx16* mutant cells is responsible for the cell migration defect that prevents cells from exiting the tailbud. We transplanted *HS:sox2* cells into the ventral margin of wild-type host embryos. Activation of *sox2* expression at bud and 12-somite stages prevented transplanted cells from exiting the tailbud into the mesodermal territory and the majority of cells are found at the posterior end of the host embryo (Fig 2A-C). To better understand the migratory dynamics of the *sox2* expressing cells in the tailbud, we labeled embryos with a nuclear localized kikume (*NLS-kikRG*), which can be photoconverted from green to red, by injecting in vitro transcribed mRNA. Small groups of cells in the NMP region were photoconverted and time-lapse imaged for 300 minutes. Wild-type cells move ventrally in a directed fashion (Fig. 2D-E, H-J). While *sox2* expressing cells move faster than wild-type cells, their overall displacement is reduced, due to significantly reduced migratory track straightness (Fig. 2F-J, F, 281 cells were tracked in 5 embryos, G, 200 cells were tracked in 3 embryos, statistics were performed using an unpaired t test ***P<0.0001). The migratory activity but lack of directed migration defines the partial EMT state during zebrafish mesoderm induction (Manning and Kimelman 2015), suggesting that *sox2* expressing cells are trapped in a partial EMT. To further confirm this, transgenic *HS:CAAX-mCherry-2A-NLS-KikGR* cells, or *HS:CAAX-mCherry-2A-NLS-KikGR* x *HS:sox2* cells, were transplanted into the ventral margin of wild-type host embryos. Sox2 expressing cells emigrated from the posterior wall of the tailbud and completed the first EMT step, but remained in the partial EMT state with dynamic membrane protrusions lacking polarization (Figure S1).

**Figure 2.**
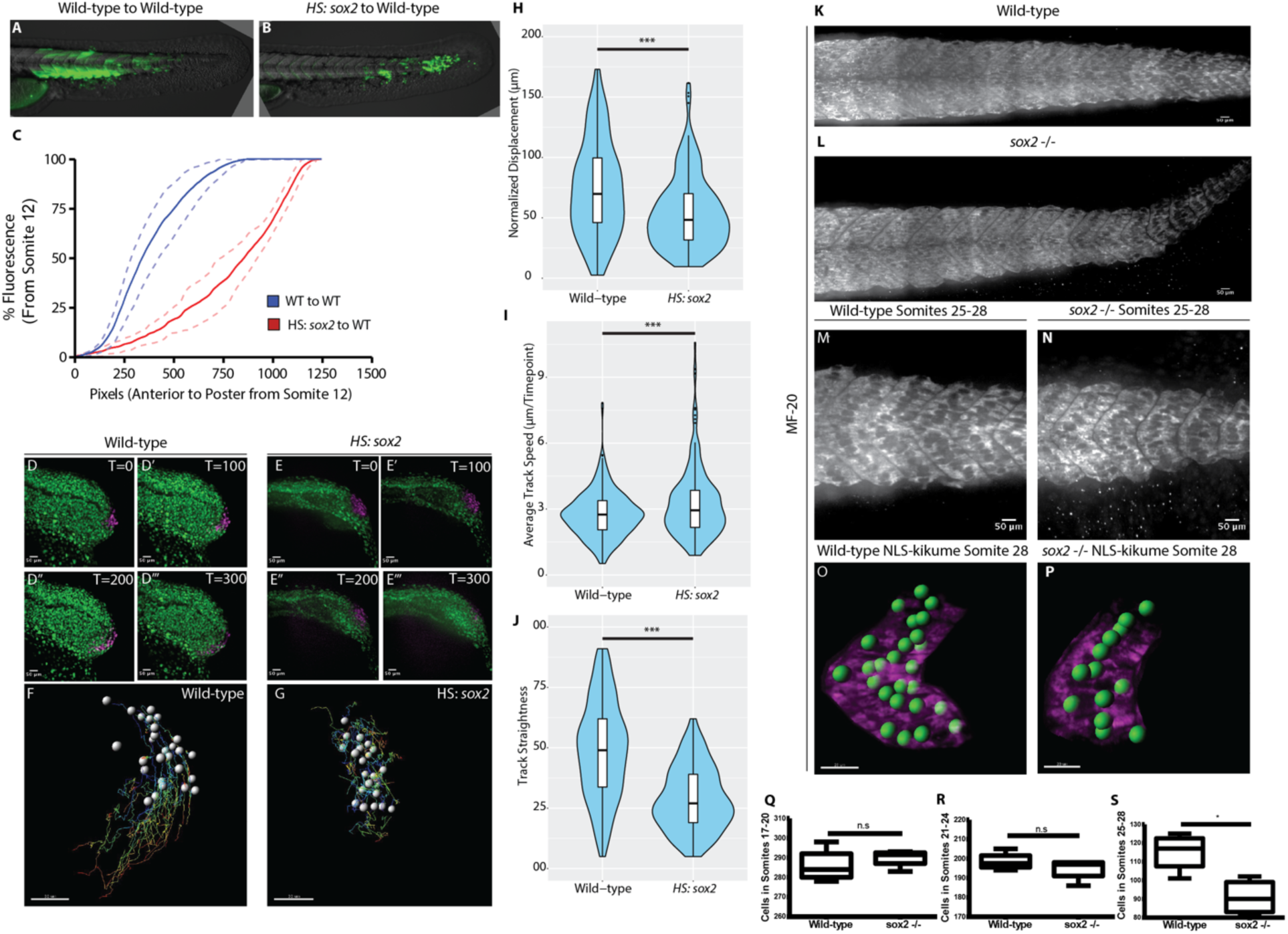
*sox2* levels control the rate of NMP exit into the mesoderm. Wild-type and *HS:sox2* embryos were injected with fluorescein dextran and cells from these embryos were transplanted into the ventral margin of shield stage wild-type host embryos (A, B, respectively). Transplants were heat shocked at bud stage and 12-somites at 39°C for 30 minutes and imaged at 36 hpf. Quantification of tailbud exit was measured as a line-scan of compound fluorescence from anterior to posterior, comparing wild-type transplanted cells (blue, N=10) with *HS:sox2* transplanted cells (red, N=6) (C). Dotted lines indicate 90% confidence. Wild-type (D-D’’’) or *HS:sox2* (E-E’’’) embryos with ubiquitous NLS-KikGR expression were photoconverted in the NMP region and time-lapse imaged for 300 minutes. Migratory tracks of photoconverted wild-type and *HS:sox2* nuclei were quantified (F, 281 cells were tracked in 5 embryos, G, 200 cells were tracked in 3 embryos, ***P<0.0001), revealing that displacement (H) and track straightness (J) were reduced in *HS:sox2* embryos, whereas average track speed was increased (I). See also Fig. S1 for analysis of cell shape in *HS:sox2* embryos. MF-20 antibody labeling of wild-type (K, M) and sox2 homozygous mutant (L, N) embryos showed that posterior somites are smaller in *sox2* mutants. Somitic nuclei were quantified, revealing that posterior somites in *sox2* mutants have significantly fewer cells than wild-type somites (O-S, Q P=0.4721, R P=0.3208, S *P=0.0145).

Since *sox2* must be repressed for NMPs to complete EMT and become fully mesenchymal, loss of *sox2* function may impact the normal rate at which cells exit the tailbud and join the paraxial mesoderm. To determine whether *sox2* function impacts the normal formation of somites from NMPs, we analyzed somite development in a *sox2* mutants (Gou et al. 2018a; Gou et al. 2018b). Somites were visualized with a muscle specific antibody (MF20, anti-myosin heavy chain), which indicated that the posterior somites appeared smaller (Fig. 2K-N). Total nuclei counts in somites 25-28 revealed that posterior somites contain significantly fewer cells than wild-type siblings (Fig. 2O-S, for panel S *P=0.0145). Our results are consistent with the hypothesis that loss of *sox2* function allows NMPs to exit into the paraxial mesoderm prematurely, leaving fewer cells to contribute to the posterior-most somites. Although we saw an increase in nuclei in more anterior somites of *sox2* mutants, these results were not statistically significant, indicating that there may be additional controls regulating somite cell number if a larger number of cells initially join a somite than normal (Fig. 2 Q P=0.4721, R P=0.3208).

### *sox2* loss of function rescues *tbx16* loss of function

Sox2 gain-of function or *tbx16* loss of function in mesoderm fated NMPs causes them to be trapped in a partial EMT state, and Tbx16 normally acts to repress *sox2* expression (Bouldin et al. 2015). To determine if *sox2* is a critical target of Tbx16 accounting for the *tbx16* mutant phenotype, we performed *tbx16* loss of function rescue experiments using a *sox2* loss of function mutant. The *sox2* mutation is able to rescue *tbx16* morphant muscle formation in both whole embryo (Fig. 3A-D’) and transplant conditions (Fig. 3E-G, 1,030 *tbx16* morphant donor cells were counted from 7 host embryos, and 609 *sox2* -/- *tbx16* morphant donor cells were counted from 4 host embryos, statistics were performed using an unpaired t test *P=0.0082), where *tbx16* morpholinos were injected into embryos from a *sox2*^+/-^ in-cross, and cells from these embryos were transplanted into wild-type host embryos. Additionally, we performed cell tracking experiments in *tbx16* MO, sox2-/- and *tbx16* MO; sox2-/- embryos (Fig. 3H-O). Cells lacking *tbx16* behave similarly to cells with a gain of *sox2* function, including decreased track straightness and overall displacement, with an increase in track speed relative to wild-type cells (Fig. 3P-R). Cells lacking both *tbx16* and *sox2* function regain wild-type like behavior, including a significant rescue of displacement, track speed, and track straightness (Fig. 3P-R, 281 wild-type cells were tracked from 5 embryos, 183 *tbx16* morphant cells were tracked from 3 embryos, 210 *sox2* -/- cells were tracked from 3 embryos, and 218 *sox2* -/- *tbx16* morphant cells were tracked from 3 embryos, statistics were performed using an unpaired t test **P=0.0029, ***P<0.0001). Taken together, these results indicate that *sox2* is a critical target gene repressed by Tbx16. In the absence of *tbx16*, increased levels of *sox2* cause cells to become trapped in a partial EMT state and prevent their exit into the mesodermal territory.

**Figure 3.**
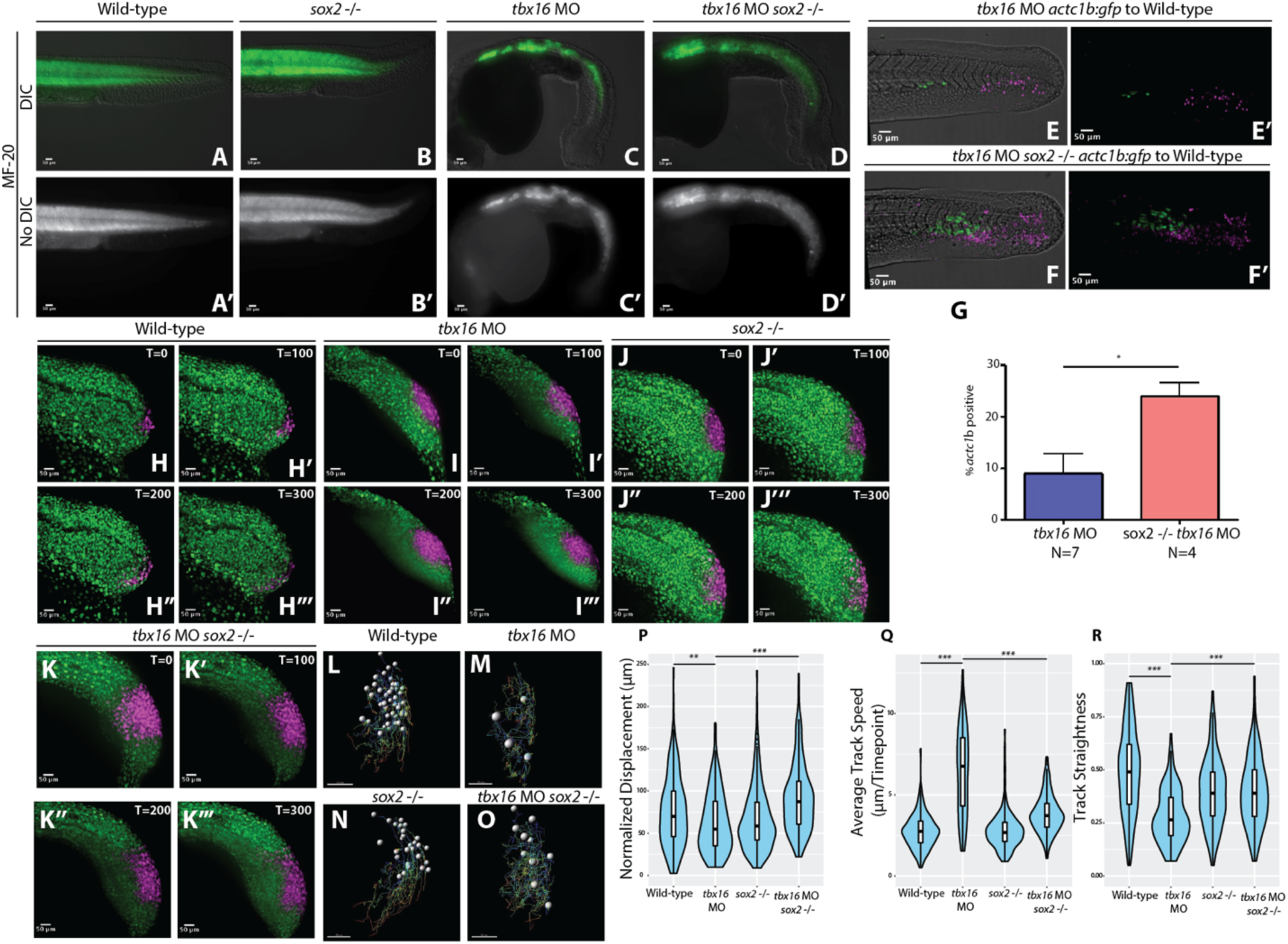
Loss of *sox2* function rescues *tbx16* loss of function. MF-20 labeling of wild-type, *sox2* -/-, *tbx16* morphant, and dual *sox2* -/- *tbx16* morphant embryos shows an increase in skeletal muscle in tbx16 morphant embryos when sox2 function is eliminated (A-D’, D, D’ compared to C, C’). Transplant experiments were performed by injecting rhodamine dextran and *tbx16* MOs into embryos from a *actc1b:gfp sox2+/-* in cross and transplanting cells into the ventral margin of wild-type host embryos. Donor cells with *sox2* function and *tbx16* loss of function showed a significantly smaller percentage of the total number of transplanted cells contributing to muscle compared to donor cells without *sox2* or *tbx16* function (E-G, 1,030 *tbx16* morphant donor cells were counted from 7 host embryos, and 609 *sox2* -/- *tbx16* morphant donor cells were counted from 4 host embryos, *P=0.0082). Statistics were performed using an unpaired T-test. N indicates number of host embryos. Wild-type (H-H’’’), *tbx16* morphant (I-I’’’), *sox2* -/- (J-J’’’), or *sox2* -/- and *tbx16* morphant (K-K’’’) embryos with ubiquitous NLS-KikGR expression were photoconverted in the NMP region and time-lapse imaged for 300 minutes. Migratory tracks of photoconverted nuclei were quantified (L-O, 281 wild-type cells were tracked from 5 embryos, 183 *tbx16* morphant cells were tracked from 3 embryos, 210 *sox2* -/- cells were tracked from 3 embryos, and 218 *sox2* -/- *tbx16* morphant cells were tracked from 3 embryos, **P=0.0029, ***P<0.0001), revealing that displacement (P), track speed (Q), and track straightness (R) were all significantly rescued towards wild-type levels in dual *sox2* and *tbx16* loss of function embryos compared to *tbx16* morphants alone.

### Checkpoint activation occurs through a synergistic interaction of Sox2 and canonical Wnt signaling

The expression of *sox2* prevents mesoderm fated NMPs from exiting the tailbud into the mesodermal territory. However, *sox2* expression does not prevent exit of NMPs from the tailbud into the spinal cord territory, suggesting that there is local difference in the niche context of the tailbud that accounts for this differential activity of Sox2. We previously showed that in the absence of canonical Wnt signaling, NMPs sustain *sox2* expression and join the spinal cord and not the mesoderm, whereas the activation of Wnt signaling using a constitutively active *β-catenin* transgene causes NMPs to join the mesoderm and not the spinal cord (Martin and Kimelman 2012). These results suggest that the presence or absence of the canonical Wnt signaling pathway accounts for the context dependent activity of *sox2*. To test this model, we performed transplant experiments with *tbx16* morphant cells or *HS:sox2* transgenic cells in the presence or absence of the *HS:TCFΔC* transgene, which cell-autonomously inhibits canonical Wnt signaling (Martin and Kimelman 2012). Transplanted wild-type cells contribute to various tissues throughout the body (Fig. 4A, N=16)). Cells lacking *tbx16* fail to join the paraxial mesoderm and instead contribute predominantly to fin mesenchyme, as previously reported (Fig. 4D-D’’, N=35) (Ho and Kane 1990; Row et al. 2011). When *sox2* or *TCFΔC* expression are activated in transplanted cells at bud stage, fewer cells contribute to the paraxial mesoderm (Fig. 4B, C, for B N=18, for C N=4). When Wnt signaling is inhibited in *tbx16* morphant cells, cells can now enter into the paraxial mesodermal territory, but rather than give rise to mesoderm, they form an ectopic spinal cord (Fig. 4E-E’’, N=43, 35 with ectopic spinal cords). The ectopic spinal cords have the proper anatomical structure of a neural canal with motile cilia projecting into the canal (Supplemental movies 1-3), as well as differentiated neurons sending axonal projections through the ectopic spinal cord (Fig. 4G-H’). To determine whether this phenotype is due to sustained *sox2* expression in *tbx16* morphant cells, we performed transplants with cells with both the *HS:sox2* and *HS:TCFΔC* transgenes. Combined heat-shock activation of *sox2* and inhibition of Wnt signaling causes the same, yet more severe phenotype of an ectopic spinal cord in the mesodermal territory along the body axis (Fig. 4F-F’’, N=17, all with ectopic spinal cords). The synergistic neural inducing activity of *sox2* activation and canonical Wnt signaling inhibition is also observed in whole embryos, where combined *sox2* activation and Wnt inhibition induces spinal cord broadly throughout the normal paraxial mesoderm domain (Fig. S2). These results show that the checkpoint holding *sox2* expressing cells in the partial EMT state is activated by the combined presence of *sox2* and canonical Wnt signaling, and that the checkpoint can be bypassed by eliminating Wnt signaling in *sox2* expressing cells, which allows them to exit the tailbud to form an ectopic spinal cord (Fig. 4I).

**Figure 4.**
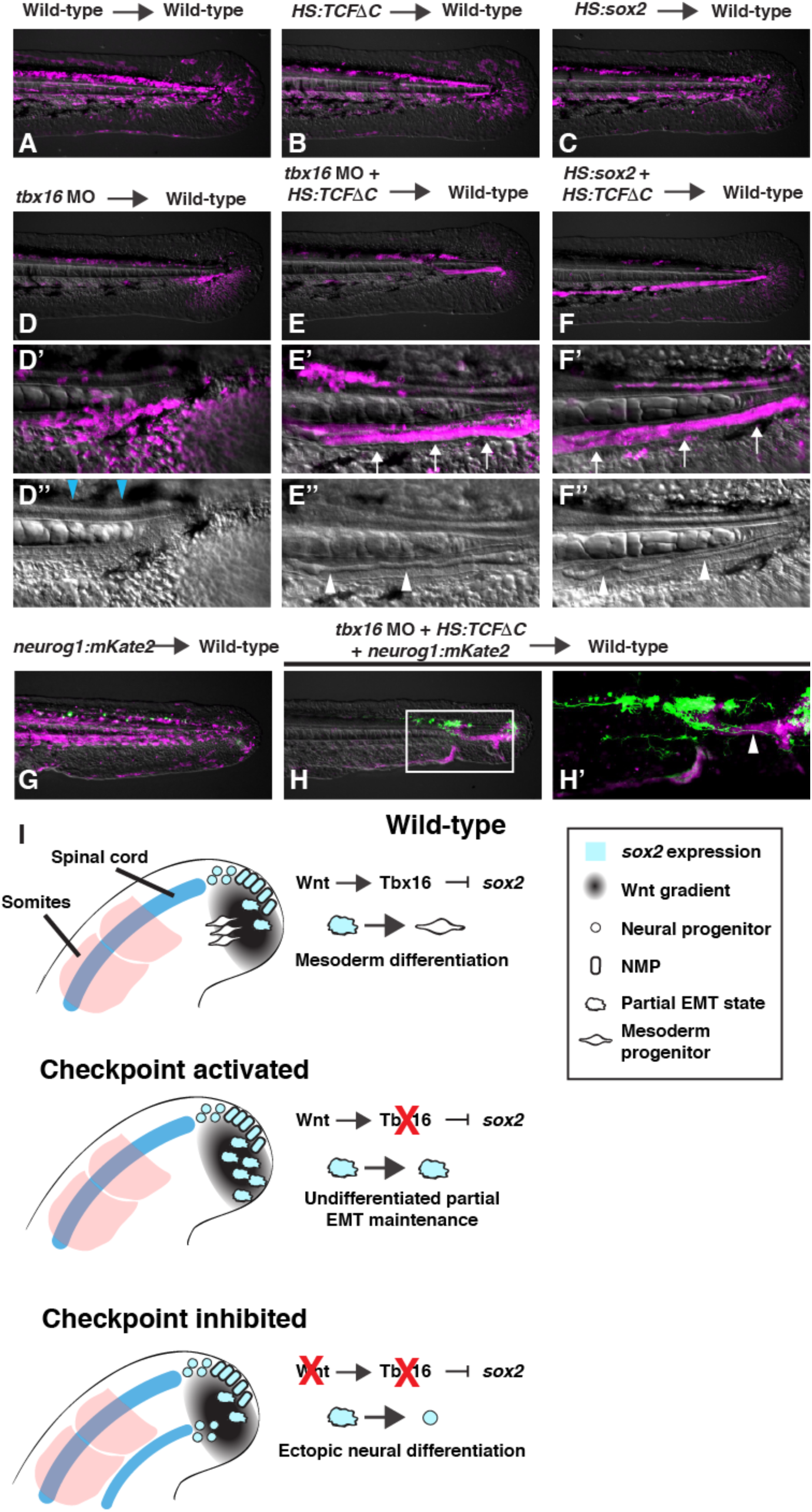
*sox2* activation in the absence of Wnt signaling results in ectopic spinal cords in transplanted cells. (A) Wild-type to wild-type transplant (N=16). (B) *HS:TCFΔC* to wild-type transplant (N=18). (C) *HS:sox2* to wild-type transplant (N=4). (D-D’’) *tbx16* mo to wild-type transplant (N=35). (E-E’’) *HS:TCFΔC tbx16* MO to wild-type transplant (N=43). (F-F’’) HS: *sox2* x *HS:TCFΔC* transplant (N=17). All transplants were performed by injecting donor embryos with 2% fluorescein dextran (false colored magenta) and transferring donor cells to the margin of 30% epiboly wild-type host embryos. All transplants were heat shocked at 40°C for 30 minutes. Loss of *tbx16* function causes donor cells that would normally form paraxial mesoderm to become fin mesenchyme (D’, blue arrowheads indicate the spinal cord, see also supplemental movie 1). Donor *tbx16* morphant cells in which Wnt signaling has been inhibited can exit the tailbud into the paraxial mesoderm territory (E’, arrows), where they form an ectopic spinal cord with a neural canal (E’’, arrowheads, see also supplemental movie 2). The same phenomenon occurs when *sox2* is activated and Wnt signaling is inhibited, where transplanted cells leave the tailbud to form an ectopic spinal cord (F’, arrows) with a neural canal (F’’, arrowheads, see also supplemental movie 3). Ectopic spinal cords formed from the combined loss of tbx16 function and Wnt signaling have differentiated neurons (green) that form long axonal projections as revealed by the *neurog1:mKate2* transgene (H, H’, arrowhead, compared to control G). See also Fig. S2 for analysis of *neurog1:mKate2* in whole embryos with loss of Wnt signaling and gain of Sox2 function. A model shows the normal progression of events as NMPs transition to paraxial mesoderm, as well as the conditions causing activation of the checkpoint (tbx16 loss of function) or checkpoint inhibited in which ectopic spinal cords form (I).

## Discussion

The mesodermal EMT during development is associated with progression towards differentiation, whereas cancer EMTs are generally thought to lead to increased stem cell characteristics and a lack of differentiation. Recent evidence suggests that metastasizing cancer cells are predominantly in a partial EMT state, and that the partial state is more stem-cell like than the fully mesenchymal state (Campbell 2018; Aiello and Kang 2019). Here we show that the partial EMT state during mesoderm induction is a developmental checkpoint that prevents differentiation into either neural or mesodermal fates. In addition to preventing differentiation, activation of the checkpoint alters the normal migratory properties of these cells. Thus, the initiation of metastasis in solid tumors through a partial EMT may be recapitulating a developmental state in which cells with aberrant gene expression patterns are activating a developmental checkpoint. Importantly, our results show that the initiation of the EMT leading to the partial EMT state is uncoupled from mesodermal fate, and is fully reversible back to the epithelial state and eventual neural differentiation by withdrawing the checkpoint activating Wnt signal. The uncoupling of EMT initiation and mesodermal fate acquisition underscores the importance of having a developmental checkpoint

Loss of *tbx16* function in zebrafish activates the developmental checkpoint because cells maintain *sox2* expression in a high canonical Wnt signaling environment. While a partial EMT state during mouse mesoderm induction has not been described, the same checkpoint is likely to function in mouse embryos, as loss of function of the closely related t-box transcription factor *Tbx6* causes a large accumulation of cells in the tailbud that are unable to exit into the mesodermal territory (Chapman and Papaioannou 1998). In this context, *sox2* also fails to be repressed and is maintained in a high Wnt environment (Takemoto et al. 2011). One key difference of the mouse *Tbx6* mutant compared to the zebrafish *tbx16* mutant is that in the mouse a subset of cells exit the tailbud to form ectopic spinal cords where somites should normally form (Chapman and Papaioannou 1998). Ectopic neural tissue is never observed in the zebrafish *tbx16* mutant, or in the *tbx16/msgn1* or *tbx16/tbx6l* double mutants, which have a more severe phenotype than the *tbx16* single mutant (Fior et al. 2012; Yabe and Takada 2012; Morrow et al. 2017). Since lowering Wnt signaling in *tbx16* mutant cells allows them to exit the tailbud and form an ectopic spinal cord, the relative level of canonical Wnt signaling inducing mesoderm in the tailbud of zebrafish is likely to be higher than in mouse. Canonical Wnt signaling promotes an accelerated exit of mesoderm from the tailbud (Martin and Kimelman 2012; Bouldin et al. 2015), and differences in Wnt signaling levels may explain the rapid exit and differentiation of mesoderm from the tailbud of zebrafish, which occurs over the course of just 12 hours, compared to the relatively slow exit from the mouse tailbud, which is drawn out over several days. Thus, modulation of Wnt signaling levels may be a key evolutionary adaptation affecting vertebrate body axis formation through changes in NMP dynamics, and the relative difference in Wnt levels may explain the species specific differences between NMP development (Martin and Kimelman 2009; Steventon et al. 2016; Attardi et al. 2018; Mallo 2019), as well as phenotypic differences of the Tbx6 mouse mutant and the *tbx16* single or *tbx16*/*tbx6l* and *tbx16*/*msgn1* double zebrafish mutants (Chapman and Papaioannou 1998; Fior et al. 2012; Yabe and Takada 2012; Morrow et al. 2017).

The developmental checkpoint preventing *sox2* positive cells from exiting into the mesodermal territory is activated by canonical Wnt signaling, and together these factors both prevent differentiation and delay morphogenesis of mesoderm fated NMPs. This is in stark contrast to the roles of these factors in the absence of the other, where each promotes differentiation along the neural (*sox2*) or mesodermal (Wnt) lineages (Takemoto et al. 2011; Martin and Kimelman 2012; Gouti et al. 2014; Garriock et al. 2015; Row et al. 2016; Gouti et al. 2017; Koch et al. 2017). This type of interaction where two lineage promoting factors can together prevent the differentiation down either lineage is a well-known feature of hematopoietic stem cells (Cross and Enver 1997; Nimmo et al. 2015). These cells are said to be in a lineage primed state, where they are held in an undifferentiated state but are poised to rapidly differentiate into either lineage as soon as one factor becomes enriched relative to the other. Our work shows that NMPs, which express *sox2* and have canonical Wnt signaling activity, are similarly in a poised state, ready to rapidly differentiate into either neural tissue or mesoderm when Wnt signaling or *sox2* expression is repressed. These results help explain the dual paradoxical functions of Wnt signaling during NMP maintenance and differentiation. While Wnt signaling is required for mesoderm induction from NMPs, it is also required for the maintenance, and possible expansion, of the undifferentiated NMP population (Takada et al. 1994; Garriock et al. 2015; Wymeersch et al. 2016). How the combination of Sox2 and Wnt signaling promotes differential cell biology than either factor alone remains to be determined, but there are several instances reported of Sox transcription factors binding to β-catenin, which in some cases can affect a unique transcriptional program (Kormish et al. 2010; Ye et al. 2014). Our results suggest the difference in Wnt function may be due to whether β-catenin is interacting predominantly with Sox2 to promote NMP maintenance, or Lef1/TCF family proteins to promote mesodermal differentiation.

## Methods

### Fish Care and Lines

All zebrafish methods were approved by the Stony Brook University Institutional Animal Care and Use Committee. Transgenic and mutant lines used include *hsp70l:sox2-2A-NLS-KikGR*^sbu100^ (referred to here as *HS:sox2*) (Row et al. 2016), *HS:CAAX-mCherry-2A-NLS-KikGR*^sbu104^ (Goto et al. 2017), *sox2-2A-sfGFP*^stl84^ (Shin et al. 2014), *actc1b:gfp*^zf13^ (Higashijima et al. 1997), *neurog1:mKate2-CAAX* (this paper), *tbx16*^b104^ (Kimmel et al. 1989), and *sox2*^x50^ (Gou et al. 2018a; Gou et al. 2018b). Heat shock inductions were performed by immersing embryos in an elevated temperature water bath (37°C to 40°C) for 30 minutes.

### Generation of a zebrafish *neurogenin1* transgenic reporter line

For the *neurog1:mKate2-CAAX* transgene, we cloned a genomic fragment spanning 8.4 kb up-stream of the *neurog1* start codon (Blader et al. 2003) into the p5E plasmid (*p5E-neurog1*, Invitrogen, USA) and the coding sequence of the fluorescent protein mKate2 (Evrogen, Russia) followed in-frame by a CAAX box from HRAS into the pME plasmid (*pME-mKate2-CAAX*, Invitrogen, USA). Using gateway recombination (Invitrogen, USA), we fused the *neurog1* genomic fragment from the *p5E-neurog1* plasmid to *mKate2-CAAX* from *pME-mKate2-CAAX* plasmid followed by a *SV40pA* signal from the *p3E-polyA* plasmid into the *pDestTol2pA2* plasmid (Kwan et al. 2007). The resultant plasmid is called *pDest-neurog1:mKate2-CAAX-SV40pA*. For transgenesis, 25 ng/μl of the *pDest-neurog1:mKate2-CAAX-SV40pA* plasmid was co-injected with *in vitro* transcribed *tol2* transposase mRNA (Thermo Fisher, USA) into one-cell-stage embryos (Kawakami et al. 2000). Transgenic fish were identified by mKate2 fluorescence at 1 dpf using a Leica M165 fluorescent stereo microscope (Leica Microsystems Inc., Germany). The full name of this transgenic line is *Tg(−8.4neurog1:Kate2-CAAX).*

### Imaging

For tailbud exit transplantation experiments, cells were mounted in 2% methylcellulose with tricaine. Imaging was done on a Leica DMI6000B inverted microscope. For cell tracking, transplant quantification, and cell shape analysis, embryos were mounted in 1% low melt agarose with tricaine and imaged on a custom built spinning disk confocal microscope with a Zeiss Imager A.2 frame, a Borealis modified Yokogawa CSU-10 spinning disc, ASI 150uM piezo stage controlled by an MS2000, an ASI filter wheel, a Hamamatsu ImageEM x2 EMCCD camera (Hamamatsu C9100-23B), and a 63× 1.0NA water immersion lens. This microscope is controlled with Metamorph microscope control software (V7.10.2.240 Molecular Devices), with laser illumination via a Vortran laser merge controlled by a custom Measurement Computing Microcontroller integrated by Nobska Imaging. Laser power levels were set in Vortran’s Stradus VersaLase 8 software.

### *In Situ* Hybridization and Immunohistochemistry

Whole-mount *in situ* hybridization was performed as previously described (Griffin et al. 1995). For skeletal muscle antibody labeling, embryos were treated with a 1:50 dilution of the MF-20 antibody (Developmental Studies Hybridoma Bank – a myosin heavy chain antibody labeling skeletal and cardiac muscle) followed by an Alexa Fluor 561-conjugated anti-mouse secondary antibody. For somite quantification embryos were injected with 100pg *kikume* mRNA and fixed at 36 hpf and treated with MF-20 antibody. MF-20-labled somites were imaged on a spinning disk confocal microscope using a 40x/1.0 dip objective in embryo media. Somitic nuclei were counted using spots on Imaris software (Bitplane, Oxford Instruments).

### Whole Embryo Reporter Expression

Reporter lines for neural (ngn:mKate2) or muscle (*actc1b:gfp*^zf13^) were crossed to the HS: *hsp70l:sox2-2A-NLS-KikGR*^sbu100^ and imaged live on a spinning disk confocal microscope using the 10x/0.3 air objective at 36 hpf.

### *sox2* Overexpression Transplants

Cell transplantation experiments were performed from sphere stage to shield stage targeting the ventral margin (Martin and Kimelman 2012). Transplanted embryos were heat shocked at 39°C for 30 minutes at bud and 12 somite stages. Embryos were imaged on the spinning disk confocal using the 10x/0.3 air objective at 36 hpf.

### *sox2* Mutant and *sox2:sfGFP* Transplants

Donor embryos were injected with 100 pg of *kikume* mRNA and a mix of two *tbx16* morpholinos (MO1: AGCCTGCATTATTTAGCCTTCTCTA (1.5ng) MO2: GATGTCCTCTAAAAGAAAATGTCAG (0.75ng)) as previously described (Lewis and Eisen 2004). Donor cells were transplanted from sphere stage donors to shield stage hosts targeted to the ventral margin as previously described (Martin and Kimelman 2012). Donor embryos were screened for the *sox2* genotype and presence of *actc1b:gfp*^zf13^ reporter. Embryos were imaged on the spinning disk confocal using the 10x/0.3 air objective at 36 hpf.

### Transplant Tissue Contribution Quantification

For neural quantification embryos were imaged live on a spinning disk confocal microscope using the 10x/0.3 air objective. For muscle quantification transplanted cells were photoconverted on an inverted microscope using 405 nm light for 30 seconds and embryos were imaged live on a spinning disk confocal microscope using the 10x/0.3 air objective. Transplanted nuclei within reporter lines were quantified using Imaris (Bitplane, Oxford Instruments). When necessary, images were stitched using Fiji (Preibisch et al. 2009).

### Transplant Cell Exit Quantification

Donor embryos were in injected with 2% fluorescein dextran and cells were transplanted from sphere to shield stage targeting the ventral margin as described previously. Host embryos were imaged on an inverted Leica DMI6000B microscope using the 10x/0.4 dry objective. Compound fluorescence from transplanted cells was measured from anterior to posterior starting from somite 12 to the end of the tail using Fiji.

### Tailbud Cell Tracking

Embryos were injected with 25 pg of *kikume* mRNA at the 1-cell stage. A small region containing NMPs was photoconverted on an inverted Leica DMI6000B microscope using 405 nm filter set for 30 seconds and tracked on a spinning disk confocal using the 20x/0.8 air objective and tracked on Imaris as previously described (Goto et al. 2017).

### Cell Shape Analysis

Cells from *HS:CAAX-mCherry-2A-NLS-KikGR*^sbu104^ donor embryos were transplanted to the ventral margin of unlabeled wild-type host embryos and imaged over 8 hours with 5 minute intervals on a spinning disk confocal microscope using the 40x/1.0 dip objective in embryo media and analyzed on Fiji and Imaris as previously described (Goto et al. 2017).

## Acknowledgments

We thank David Matus for use of the spinning disc confocal microscope and comments on the manuscript, Taylor Kinney for advice, Neal Bhattacharji and Stephanie Flanagan for excellent zebrafish care, and Bruce Riley for providing sending the *sox2*^*x50*^ mutant prior to publication. This work was supported NIH NINDS (R01NS102322) to HK, and by NSF (IOS 1452928) and NIH NIGMS (1R01GM124282) grants to BLM.

---

**Figure S1 (related to Figure 2):**
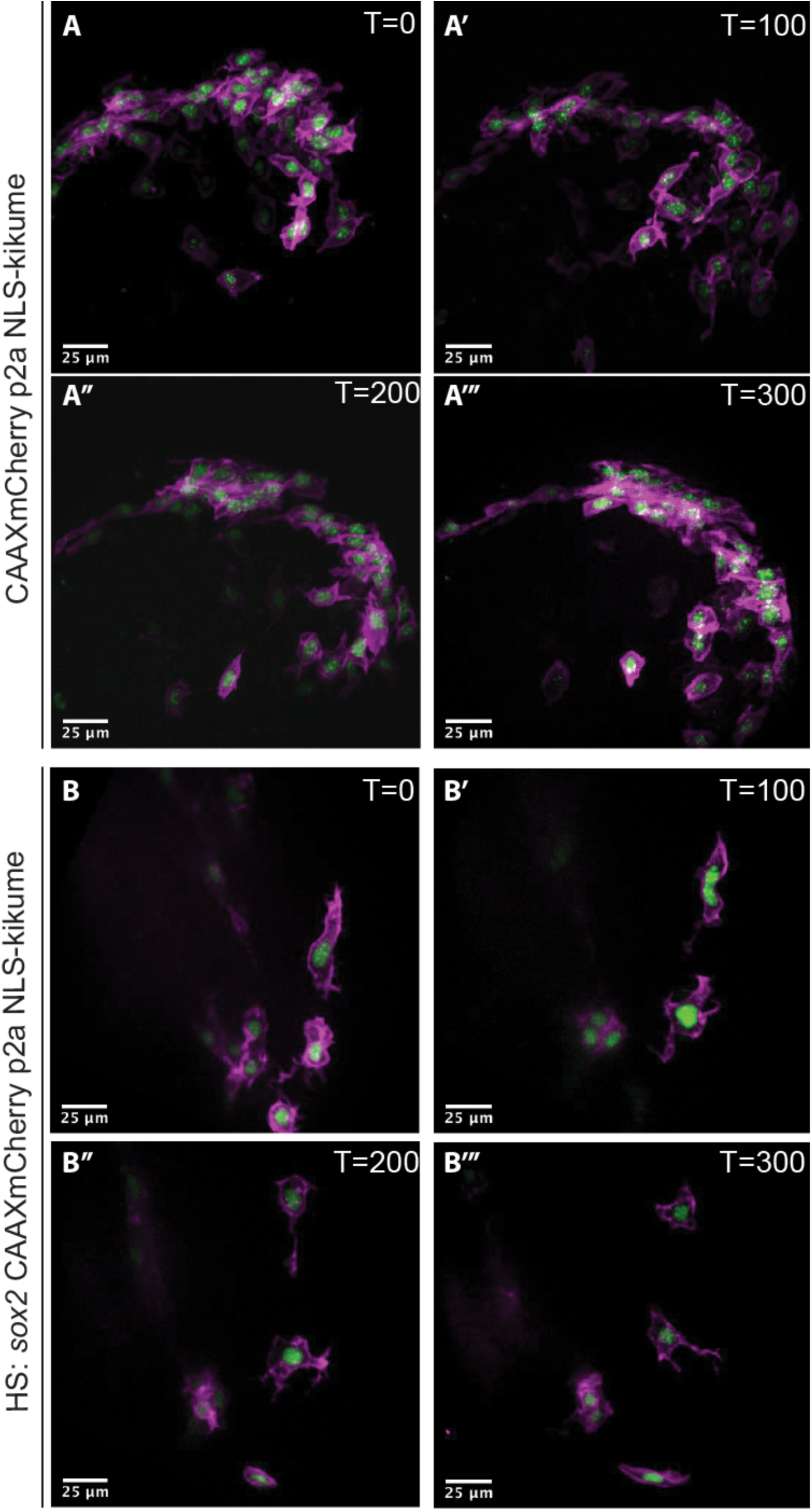
Sox2 gain of function does not prevent the first EMT step of NMPs during mesoderm induction. Donor cells from *HS:CAAX-mCherry-2A-NLS-KikGR* (A-A’’’) or *HS:CAAX-mCherry-2A-NLS-KikGR* + *HS:sox2* (B-B’’’) embryos were transplanted into the ventral margin of shield stage wild-type host embryos and heat-shocked at bud stage and 12-somites at 39°C for 30 minutes. Transplanted cells in which Wnt signaling is inhibited fail to undergo the first step of EMT and remain in the posterior wall NMP epithelium. Cells with sox2 gain of function on the other hand are able to complete the first EMT step as indicated by their protrusive activity but do not complete the second EMT step (B-B’’’).

**Figure S2 (related to Figure 4).**
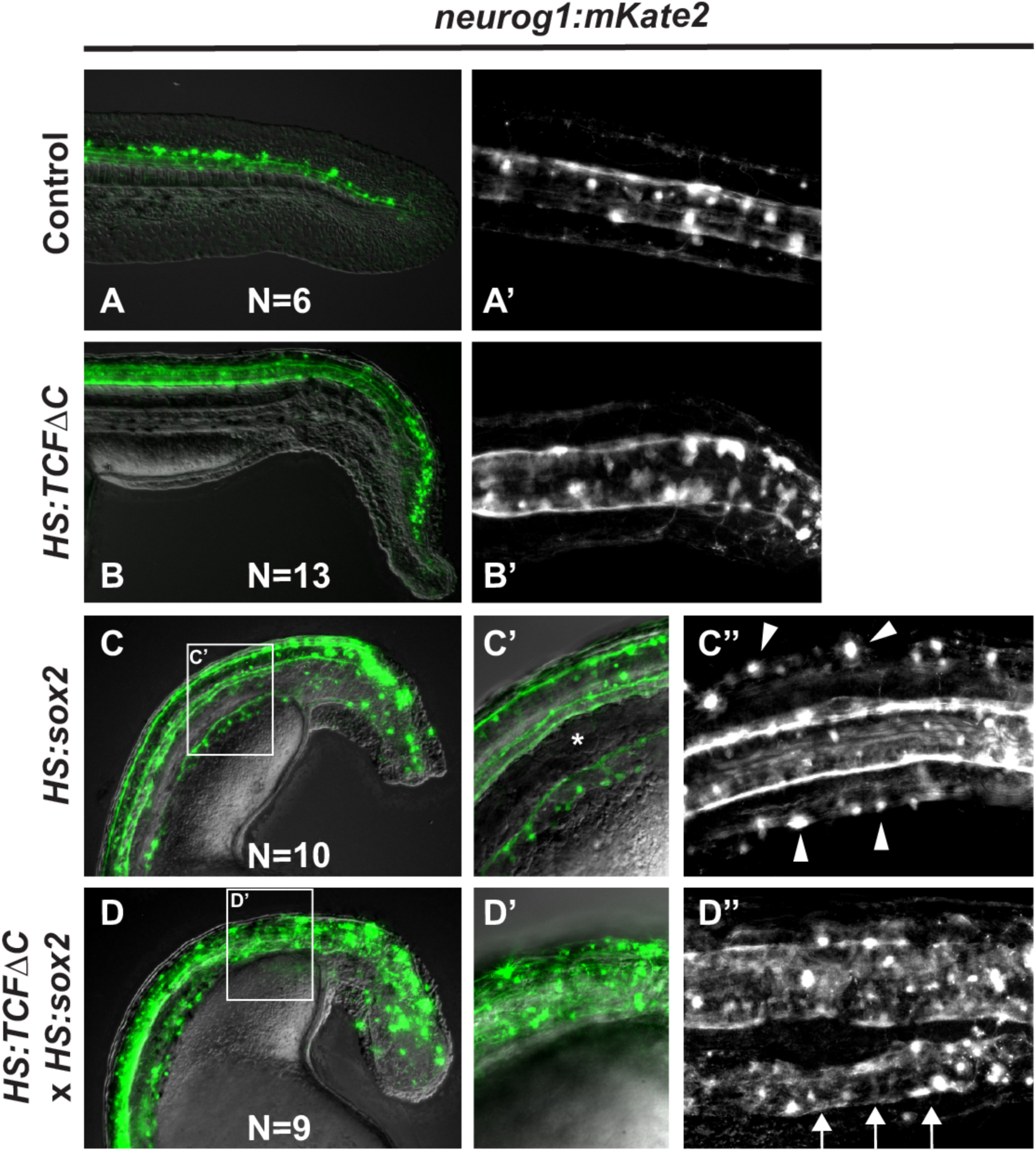
Synergy between gain of *sox2* function and loss of Wnt function during NMP transition to mesoderm. The *neurog1:mkate2* reporter line to monitor neural fate in wild-type (A, A’), Wnt loss of function (B, B’), sox2 gain of function (C-C’’), or combined sox2 gain of function and Wnt loss of function (D-D’’). Embryos were heat-shocked at bud stage and imaged at 48 hpf from a lateral view (A, B, C, C’, D, D’) or a dorsal view (A’, B’, C’’, D’’) with anterior to the left. Control (A, A’) and Wnt loss of function (B, B’) embryos never show ectopic neural tissue. Sox2 gain of function embryos have ectopic neurons in the ventral paraxial region, which are separated from the spinal cord by small somites (C’, star). The neuron cell bodies are present in lateral positions to the spinal cord (C’’, arrowheads). The combined loss of Wnt function and gain of sox2 function causes a robust expansion of spinal cord fate in the paraxial mesoderm territory (D’ compared to C’), which when visualized from a dorsal view shows the existence of an ectopic spinal cord in the paraxial mesoderm territory (D’’, arrows).

**Supplemental movie 1 (related to Figure 4)-DIC movie of a wild-type host embryo with *tbx16* MO donor cells (the same embryo pictured in Figure 4D-D’’).** Motile cilia can be observed beating in the neural canal of the spinal cord (spinal cord is indicated by blue arrowheads in Figure 4D’’).

**Supplemental movie 2 (related to Figure 4)-DIC movie of a wild-type host embryo with *tbx16* MO + *HS:TCFΔC* donor cells (the same embryo pictured in Figure 4E-E’’).** Motile cilia can be observed beating in the neural canal of the ectopic spinal cord generated by transplanted cells (ectopic spinal cord is indicated by white arrowheads in Figure 4E’’).

**Supplemental movie 3 (related to Figure 4)-DIC movie of a wild-type host embryo with *HS:sox2* + *HS:TCFΔC* donor cells (the same embryo pictured in Figure 4F-F’’).** Motile cilia can be observed beating in the neural canal of the ectopic spinal cord generated by transplanted cells (ectopic spinal cord is indicated by white arrowheads in Figure 4F’’).

